# Behavioral and postural analyses establish sleep-like states for mosquitoes that can impact host landing and blood feeding

**DOI:** 10.1101/2021.11.11.467918

**Authors:** Oluwaseun M. Ajayi, Justin M. Marlman, Lucas A. Gleitz, Evan S. Smith, Benjamin D. Piller, Justyna A. Krupa, Clément Vinauger, Joshua B. Benoit

## Abstract

Sleep is an evolutionarily conserved process that has been described in different animal systems. For insects, sleep characterization has been primarily achieved using behavioral and electrophysiological correlates in a few systems. Sleep in mosquitoes, which are important vectors of disease-causing pathogens, has not been directly examined. This is surprising as circadian rhythms, which have been well studied in mosquitoes, influence sleep in other systems. In this study, we characterized sleep in mosquitoes using body posture analysis and behavioral correlates, and quantified the effect of sleep deprivation on sleep rebound, host landing and blood-feeding propensity. Body and appendage position metrics revealed a clear distinction between the posture of mosquitoes in their putative sleep and awake states for multiple species, which correlate with a reduction in responsiveness to host cues. Sleep assessment informed by these posture analyses indicated significantly more sleep during periods of low activity. Nighttime and daytime sleep deprivation resulting from the delivery of vibration stimuli induced sleep rebound in the subsequent phase in day and night active mosquitoes, respectively. Lastly, sleep deprivation suppressed host landing in both laboratory and field settings and also impaired blood feeding of a human host when mosquitoes would normally be active. These results suggest that quantifiable sleep states occur in mosquitoes, and highlight the potential epidemiological importance of mosquito sleep.

## Introduction

Sleep is a phenomenon universally observed across the animal kingdom with notable description in cnidarians [1], nematodes [2], arthropods [3, 4], and mammals [5]. During sleep, animals lose connection with their external environment, as a result of attenuated sensory processing and motor outputs which pose significant predation risks to the individuals [6]. In this process, individuals cannot search for food resources, engage in parental care or evade detrimental situations, which indicate that sleep is of essential benefits when considering its trade-offs [4]. In vertebrates (particularly mammals), acute sleep deprivation results in impaired cognition [7, 8], while chronic sleep deprivation has been implicated in hallucinations, speech delay, and sometimes death [9, 10]. Similarly, the importance of sleep has been established in invertebrates, especially in insects. Studies have shown that sleep deprivation significantly reduces the precision of waggle dance signaling in honey bees (*Apis mellifera*) [11], and results in short- and long-term memory defects, along with a multitude of other factors, in fruit flies (*Drosophila melanogaster*) [12, 13].

Characterization of parameters underlying sleep is evaluated using different approaches [5]. However, the two classical and robust hallmarks of sleep-like states in a variety of animals have been behavioral and/or electrophysiological correlates [14, 15]. Modulations in brain wave activity, which is measured using electroencephalography in mammals or recordings of local field potentials in invertebrates, can establish specific electrophysiological correlates of sleep [16, 17]. Behaviorally, sleep can be characterized using the following features: (i) species-specific postures, (ii) reversible prolonged quiescence in certain periods in the circadian day, (iii) increased arousal threshold or decreased response to stimuli, and (iv) rebound or recovery sleep in response to sleep deprivation [15]. For many animal systems, the establishment of behavioral factors is sufficient in characterizing the sleep-like state.

Despite the characterization of sleep in insect systems including fruit flies [18, 19], cockroaches [3], bees [20, 21], and wasps [22], and the likely benefits of sleep [23–25], little is known about sleep in blood-feeding arthropods. There has been limited focus on sleep in mosquitoes, unlike established roles of circadian rhythm (which is linked to/influences sleep in many animals) on mosquito biology [26–28]. The entirety of sleep-based research in mosquitoes may be restricted to only two studies: (i) an early study on the resting postures of *Aedes aegypti* [29], but this study did not consider these resting postures as sleep-like states, and (ii) our recent review that provides lines of evidence for sleep-like conditions in mosquitoes, including the potential of unique postural differences between sleep-like and awake states in a single mosquito, *Ae. aegypti* [30].

In this study, we provide the characterization of sleep-like states in mosquitoes based on behavioral features established in other systems and show the effect of sleep deprivation on epidemiologically relevant aspects of mosquito biology: their locomotor activity, host landing, and blood-feeding propensity. Our results indicate that; sleep-like states occur in mosquitoes with quantifiable postural metrics which correlate with increased arousal threshold, and mosquito sleep deprivation induces subsequent sleep rebound and impairs host landing and blood feeding during normally active periods. This first extensive evaluation in mosquitoes represents an ideal model for understanding the importance of sleep in blood feeding arthropods.

## Results

### Distinct postural differences exist between putative sleep-like and active (awake) states in multiple mosquito species

Sleep states induce a behavioral quiescence typically associated with an animal-specific stereotypical posture [31–34]. In *Ae. aegypti*, we previously showed that prolonged immobilization was associated with a prostrate state where the hind legs are lowered and the thorax and abdomen brought closer to the substrate [29]. Here, we examined whether different postural states occur across mosquito species and if these states are correlated with prolonged periods of inactivity. We video recorded adult *Ae. aegypti, Culex pipiens*, and *Anopheles stephensi* females in groups of 20 females within acrylic containers (16 oz mosquito breeder; BioQuip, Rancho Dominguez, CA, USA) whose top was covered by a fabric mesh. After an acclimatization period of 2 hours to reduce the impact of previous host manipulation, pictures were taken from outside the experimental room by remote accessing the computer controlling the camera (Figure 1A). Principal Component Analysis (PCA) of the hind leg angle relative to the mosquito’s main body axis, the body angle relative to the substrate, the elevation of the hind leg relative to the substrate, and the elevation of the thorax relative to the substrate, revealed a clear clustering of each species’ body posture (ANOSIM: R = 0.204; *p* < 0.001) and a distinct clustering of postures associated with mosquitoes in prolonged immobilization (>30 min) (ANOSIM: R = 0.824; *p* < 0.001) (Figure 1B). Interestingly, the analysis of similarity’s R statistics, which compares the mean of ranked dissimilarities between groups to the mean of ranked dissimilarities within groups, revealed a stronger dissimilarity between sleep/activity states than between mosquito species (R = 0.824 and 0.204, respectively). Analysis of the contribution of each variable to the principal components (PCs) revealed that the hind leg angle contributed to 99.4% of the variance explained by PC1 (88.9%), and the body angle contributed to 99.3% of the variance explained by PC2 (11%). In other words, while the body angle seems mostly driven by interspecific differences, the position of the hind legs appears as a reliable indicator of prolonged rest states.

**Figure 1:**
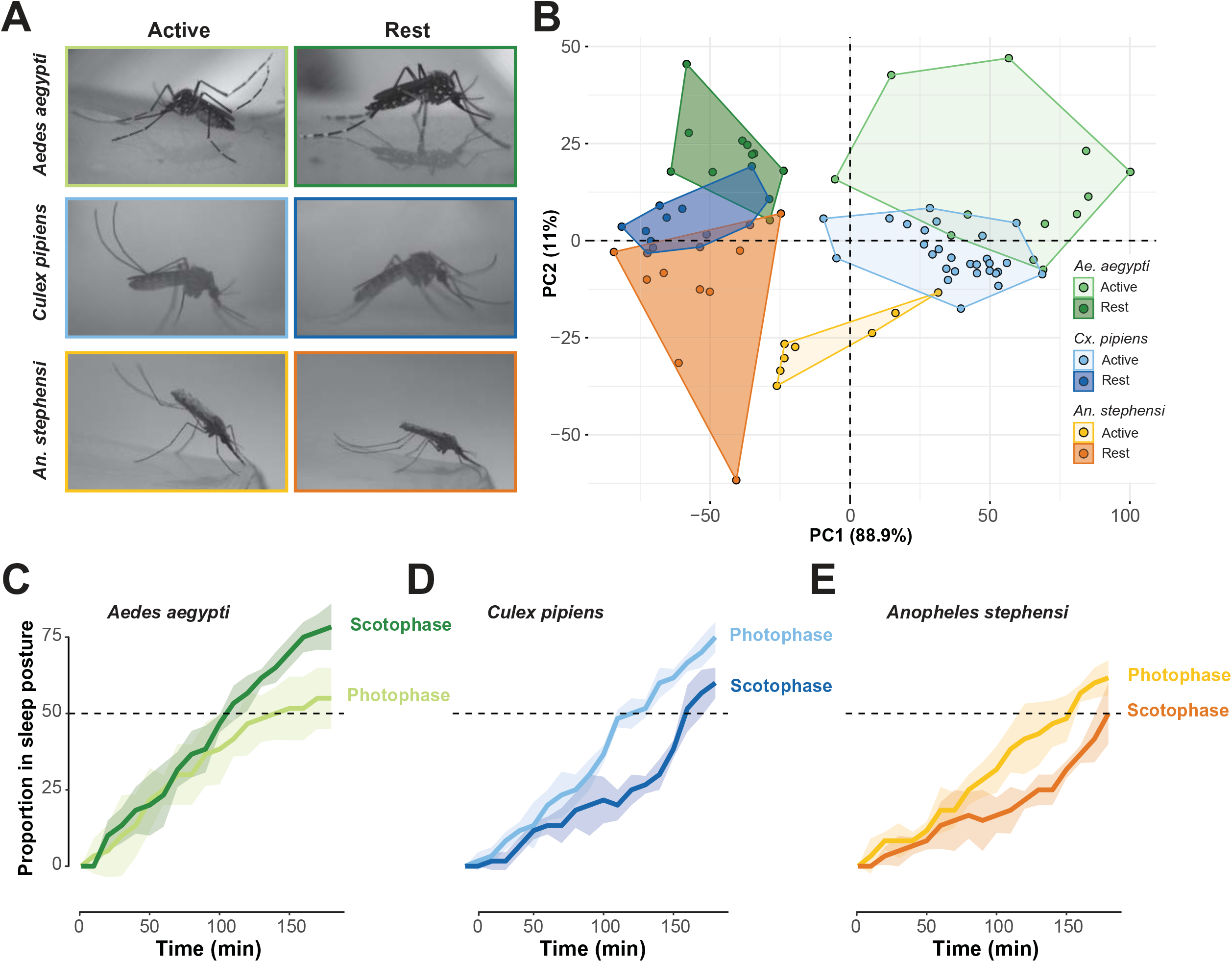
Prolonged inactivity is associated with stereotypical body postures in multiple mosquito species. (A). Representative pictures of adult females *Aedes aegypti* (top row), *Culex pipiens* (middle row), and *Anopheles stephensi* (bottom row) either in active state (left column) or at rest (right column). (B). Principal component analysis of the ensemble of postural measures. The colors of points and grouping contours indicate the species and status of each point: Green: *Ae. aegypti* (n = 22); blue: *Cx. pipiens* (n = 41); orange: *An. stephensi* (n = 17). Darker colors indicate rest and lighter colors indicate active states. (C). Proportion of mosquitoes displaying a sleep posture as a function of time for each species (n = 3 replicates; N = 120 individuals each). Proportions were quantified either during the photophase (lighter colors), or during the scotophase (darker colors).

In a second postural assay, adult females of *Ae. aegypti*, *Cx*. *pipiens*, and *An. Stephensi* were individualized in plastic *Drosophila* tubes (25mm x 95mm, Genesee Scientific, San Diego, CA, USA) and, for each assay, 20 tubes of females of the same species were positioned horizontally, in the field of view of a video camera (C920, Logitech, Lausanne, Switzerland). Every 10 minutes, the posture of each individual was recorded and classified as ‘active’ or ‘rest’ based on the angle of the hind legs relative to the main body axis. For all three species, regardless of whether the experiment was conducted during the last 3 hours of the photophase, or during the first 3 hours of the scotophase (to capture the activity peaks of both nocturnal and diurnal species), the proportion of mosquitoes in a sleep-like posture was strongly correlated with the amount of time spent in the absence of external stimulation (Figures 1C, 1D and 1E; *Ae. aegypti* photophase: Pearson correlation coefficient *r =* 0.907; scotophase: *r* = 0.978; *Cx. pipiens* photophase: *r* = 0.983; scotophase: *r* = 0.929; *An. stephensi* photophase: *r* = 0.959; scotophase: *r* = 0.906; n= 30 each). Although a log-rank test revealed no significant differences between scotophase and photophase sleep curves, the amount of time required for 50% of individuals to be in a sleep-like posture was larger during the photophase than during the scotophase for the diurnal *Ae. aegypti*, conversely to the nocturnal *Cx. pipiens* and *An. stephensi*.

Of importance is that in *Cx. pipiens* and *Ae. aegypti*, there is a reduction in response to host cues, indicated by a reduction in flight activity triggered by the presence of an experimenter, for individuals in prolonged resting/sleep state (Table 1). This provides evidence that the sleep-states are likely correlated with increased arousal threshold. Overall, these results indicate that there are distinct postures associated with putative sleep-like states in mosquitoes, that individuals will enter these postural states more rapidly during the circadian period associated with lower activity, and that these states correlate with increased arousal thresholds in both diurnal and nocturnal species.

**Table 1.** Prolonged inactive, sleep-like periods reduce the responsiveness of mosquitoes to a potential host. Data represent the percent response (mean +/- SE) that show flight within thirty seconds following exposure to experimenter breath.

| Time (minutes) of inactivity | *Aedes aegypti* | *Culex pipiens* |
| --- | --- | --- |
| 0 | 0.85 +/- 0.10<sup>a</sup> | 0.54 +/- 0.15<sup>a</sup> |
| 30 | 0.82 +/- 0.12<sup>a</sup> | 0.44 +/- 0.17<sup>a,b</sup> |
| 60 | 0.67 +/- 0.14<sup>a,b</sup> | 0.36 +/- 0.15<sup>a,b</sup> |
| 120 | 0.40 +/- 0.22<sup>b</sup> | 0.25 +/- 0.22<sup>a,b</sup> |
| 240 | 0.46 +/- 0.15<sup>b</sup> | 0.21 +/- 0.12<sup>b</sup> |

### Circadian timing and amount of sleep-like period differ among multiple mosquito species

One important hallmark of sleep is that organisms (studied so far) experience reversible prolonged periods of immobility/inactivity during a particular phase of the circadian day [2,18,19,35,36]. To determine periods of putative sleep (lack of activity) in mosquitoes, we quantified the rest-activity rhythm of *Ae. aegypti, An. stephensi,* and *Cx. pipiens* using an infrared-based activity monitoring system during a 24-hr circadian day. In *Drosophila*-based studies, sleep is usually defined as a period of inactivity lasting at least for 5 minutes and the occurrence of rest (putative sleep) is inversely related to the number of activity counts (beam breaks) recorded per a given time [18,19,37]. This short period of 5 minutes is not appropriate for mosquitoes. Rather, we quantified the sleep profile for mosquitoes using a period of inactivity lasting 120 minutes based on the time required for 50% of mosquitoes to enter a sleep-like state (Figures 1C, 1D and 1E).

Based on historical observations of field-based mosquito feeding behavior, we hypothesized that *Ae. aegypti* (a diurnal mosquito *i.e*., “day biter” [38, 39]) will have increased activity during the photophase (day time) and rest *i.e*., putative sleep will be well consolidated in the scotophase (night time). Laboratory measurements in *Ae. aegypti* show that activity increases from mid-day till the onset of light off, but activity reduces significantly throughout the night after light off (Figure 2A). Putative sleep for *Ae. aegypti* decreases from mid-day till the end of the photophase, but putative sleep is well consolidated in the scotophase (Figure 2D). This is an exact inverse of what we reported in the activity profile.

**Figure 2:**
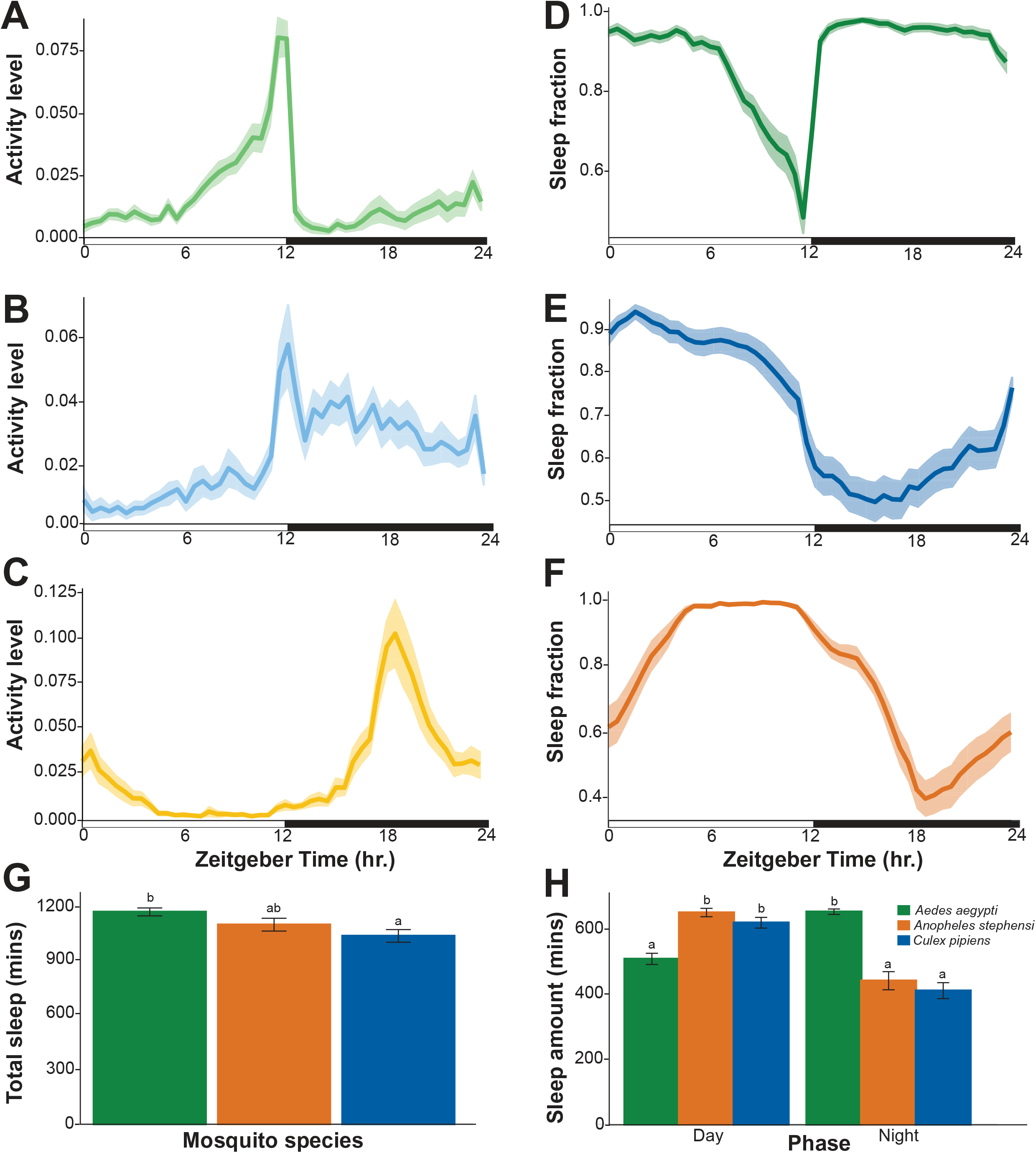
Timing and amount of sleep differ among multiple mosquito species. Basic activity rhythm of (A) *Aedes aegypti*, (B) *Culex pipiens* and (C) *Anopheles stephensi* over an 24-hour period. The y axis represents the mean beam crosses made by all the mosquitoes. Sleep profile of (D) *Aedes aegypti*, (E) *Culex pipiens* and (F) *Anopheles stephensi* averaged into a single 24-hour period. The y axis shows the proportion of time spent sleeping (defined as inactive periods of 120 minutes), averaged for each mosquito within a 30 min time window. The x axis for all the plots represents the Zeitgeber time (ZT0 - ZT24). The solid lines and the shaded areas display means and their 95% bootstrap confidence interval, respectively. White and black horizontal bars represent the photophase and scotophase, respectively. Comparison of (G) total sleep and (H) daytime and nighttime sleep among the three mosquito species. Error bars denote SE of the mean sleep amount. Different letters indicate significant differences between treatment groups (Kruskal-Wallis Test with Dunn’s multiple comparison post hoc, *p* < 0.05). In all the analyses, n = 60 for both *Aedes aegypti* and *Culex pipiens* and n = 34 for *Anopheles stephensi*.

Comparative analysis conducted in *Cx. pipiens* (a crepuscular-dark active species [40]) shows that activity is consistently low during the day but increases in anticipation of light off *i.e.*, dusk (Figure 2B) and putative sleep is reduced significantly from dusk into the first-half of the night (Figure 2E). For the nocturnal mosquito *i.e*., “night biter” [28], *An. stephensi* increased activity from early night into the mid-night, with activity reducing as day approaches (Figure 2C). Putative sleep occurred throughout the day for *An. stephensi* (Figure 2F).

Sleep amount in minutes was quantified for our laboratory strains of mosquitoes, with comparisons made among the three species. There was a significant difference in the mean total sleep among the three mosquito species (Kruskal-Wallis test: *X*^2^ = 7.221; *p* = 0.027), where the difference exists only between *Ae. aegypti* and *Cx. pipiens* (Dunn’s multiple comparison: *p* = 0.027; Figure 2G). As expected, daytime and nighttime sleep amount differs among the species (Kruskal-Wallis test: daytime, *χ*^2^ = 36.831; *p* < 0.001; nighttime, *X*^2^ = 65.519; *p* < 0.001; Figure 2H); however, there was no difference between *Cx. pipiens* and *An. stephensi* for both day and night. Together, these results reveal the marked differences in timing and amount of sleep-like periods in different mosquitoes species.

### Sleep deprivation induces sleep rebound in *Aedes aegypti* and *Anopheles stephensi* depending on the phase of perturbation

Sleep deprivation in mosquitoes was assessed for subsequent sleep rebound when individuals are normally active. In *Ae. aegypti*, sleep deprivation by mechanical disturbance was conducted for 12 hrs during the night, 4 hrs during the night, and 12 hrs during the day. However, in *An. stephensi*, sleep deprivation was only done for 12 hrs during the day for comparative observations with the day-active *Ae. aegypti*.

*Ae. aegypti* mosquitoes subjected to sleep deprivation throughout the night recorded a significant sleep loss of about 558 minutes when you compare with sleep amount of the preceding night (Wilcoxon signed rank test: V = 1122; *p* < 0.001; Figure S1E). This sleep loss promoted a significant rebound in the subsequent photophase, with a gain of approximately 76 minutes of sleep (Paired t-test: *t* = 3.463; *p* = 0.001; Figure 3A). A significant sleep loss of nearly 159 minutes occured in *Ae. aegypti* mosquitoes that experienced sleep disruption in the first four hours of the night (Wilcoxon signed rank test: V = 1672.500; *p* < 0.001; Figure S1F). Even this short amount of lost sleep early in the night was adequate to induce sleep rebound in the following day; a significant sleep gain of nearly 1 hour was reported (Paired t-test: *t* = 3.846; *p* < 0.001; Figure 3B). In the *Ae. aegypti* mosquitoes subjected to sleep deprivation during the photophase, the amount of sleep lost by comparing with sleep amount in the baseline day was approximately 436 minutes (Wilcoxon signed rank test: V = 2013; *p* < 0.001; Figure S1G). However, this failed to yield a significant sleep gain in the subsequent night, indicating that sleep deprivation during a normally active period does not generate a rebound (Paired t-test: *t* = 0.378; *p* = 0.707; Figure 3C). This was not the case for *An. stephensi* mosquitoes, as daytime sleep deprivation in this species mirrored that of nighttime sleep deprivation in *Ae. aegypti*, which was expected as this species is active at night. A significant sleep loss of about 594 minutes was reported (Wilcoxon signed rank test: V = 666; *p* < 0.001; Figure S1H), which induced a significant sleep recovery in the subsequent scotophase (Wilcoxon signed rank test: sleep gain = 196 mins; V = 73; *p* < 0.001; Figure 3D).

**Figure 3:**
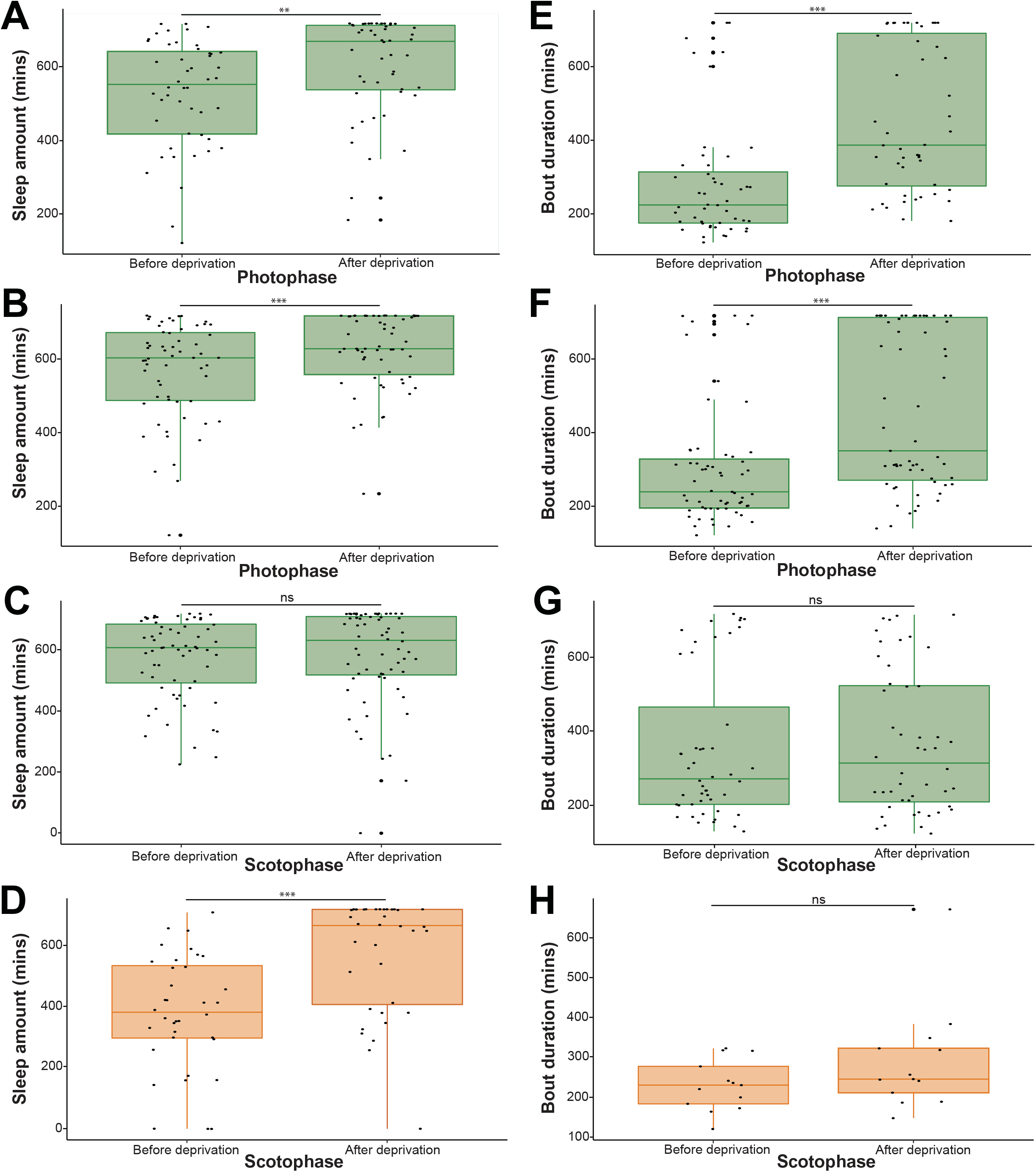
Sleep deprivation induces sleep rebound in both *Aedes aegypti* and *Anopheles stephensi*. Comparison of sleep amounts before and after sleep deprivation in (A) 12hr nighttime sleep deprivation experiment in *Aedes aegypti* (n = 48), (B) 4hr nighttime sleep deprivation experiment in *Aedes aegypti* (n = 59), (C) 12hr daytime sleep deprivation experiment in *Aedes aegypti* (n = 64) and (D) 12hr daytime sleep deprivation experiment in *Anopheles stephensi* (n = 36). Comparison of average bout durations before and after sleep deprivation in (E) 12hr nighttime sleep deprivation experiment in *Aedes aegypti* (n = 48), (F) 4hr nighttime sleep deprivation experiment in *Aedes aegypti* (n = 59), (G) 12hr daytime sleep deprivation experiment in *Aedes aegypti* (n = 48, individuals with zero values were excluded) and (H) 12hr daytime sleep deprivation experiment in *Anopheles stephensi* (n = 13, individuals with zero values were excluded). Test of significant difference between groups was carried out using paired t-test or wilcoxon signed rank test where applicable (ns = not significant, ** = *p* < 0.01, *** = *p* < 0.001).

The influence of sleep deprivation in mosquitoes was also examined in relation to another sleep architecture *i.e*., sleep bout duration. Results showed that sleep deprivation promoted a significantly increased sleep bout duration in the subsequent light phase for both the 12 hrs nighttime (Wilcoxon signed rank test: V = 994; *p* < 0.001; Figure 3E) and 4 hrs nighttime deprivations in *Ae. aegypti* (Wilcoxon signed rank test: V = 1364; *p* < 0.001; Figure 3F). As expected, daytime sleep deprivation in *Ae. aegypti* did not significantly impact sleep bout duration in the subsequent night (Paired t-test: *t* = 0.481; *p* = 0.633; Figures 3G and S1I). Although, daytime sleep deprivation in *An. stephensi* significantly promoted sleep gain in the subsequent night, sleep bout duration was not significantly affected (Wilcoxon signed rank test: V = 64; *p* = 0.216; Figures 3H and S1J). From our results, mosquitoes deprived of sleep during the normal periods of low activity, experience sleep rebound in the subsequent phase, but there was no sleep recovery if sleep deprivation occurs during their normally active period.

### Sleep deprivation in *Aedes aegypti* suppresses host landing in both laboratory and field settings, and impairs blood-feeding propensity

The impact of sleep deprivation on host landing in *Ae. aegypti* both in laboratory and field mesocosm experiments was assessed to establish a specific role in relation to interactions with potential hosts. In specific, the number of mosquitoes that landed on an artificial host 4 hours after a long-night sleep deprivation were assessed at different time points. In the laboratory assay, the proportion of mosquitoes that landed was lower in the sleep deprived group when compared with control at all time points (Figure 4A). Similar results were also observed in the field, with a lesser proportion of mosquitoes landing on the artificial host at all time points in the sleep deprived group in comparison with the control counterparts (Figure 4B).

**Figure 4:**
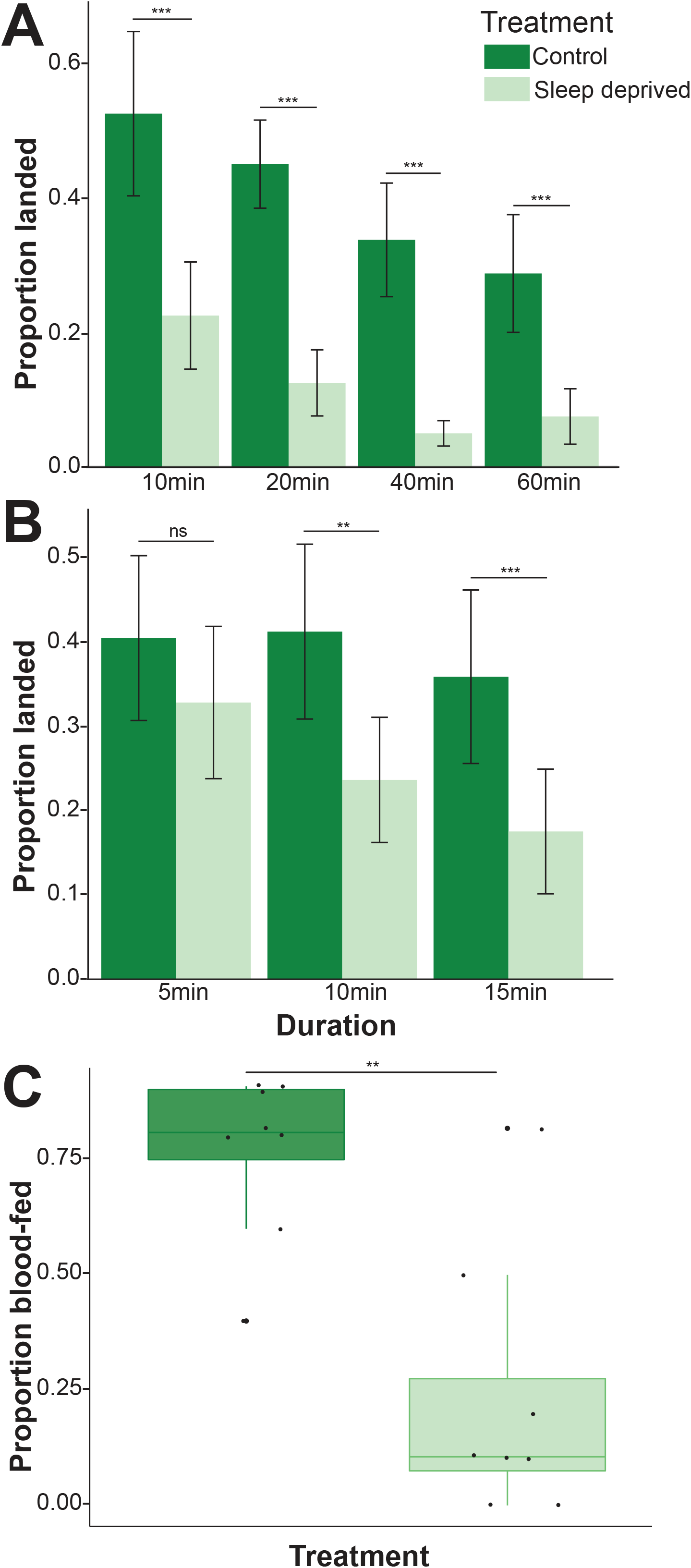
Host landing and blood-feeding propensity are impaired by sleep deprivation in *Aedes aegypti*. Mean proportion of *Aedes aegypti* mosquitoes that landed on the artificial host at different time points following sleep deprivation during the subsequent photophase in the (A) laboratory assay (n = 8 tests of 10 mosquitoes each) and (B) field mesocosm experiment (n = 13 tests of 10 mosquitoes each). (C) Proportion of *Aedes aegypti* mosquitoes that blood fed during the subsequent photophase after sleep deprivation (n = 8 tests of 10 mosquitoes each). Error bars denote SE of the mean proportion of mosquitoes that landed on the artificial host. A general linear model and wilcoxon rank sum test were used to assess significant differences in host landing and blood feeding between the treatment groups, respectively (ns = not significant, ** = *p* < 0.01, *** = *p* < 0.001).

A general linear model assessing host landing status (‘landed’ and ‘not landed’) relative to treatment (sleep deprived and control) was utilized to examine for significance. Host landing was significantly explained by sleep deprivation in the lab-based assay (*p* < 0.001 for all time points, Figure 4A). In the field-based studies (Figure 4B), no significance was noted at 5 mins (*p* = 0.199) but variation in host landing was significantly explained by sleep deprivation at 10 mins (*p* = 0.002) and 15 mins (*p* < 0.001).

In addition, we evaluated the effect of sleep deprivation on blood-feeding propensity, as a proxy for vectorial capacity. This was done by quantifying the number of mosquitoes that blood fed on a volunteer host, 4 hours after a 12-hr nighttime sleep deprivation. Result shows that sleep deprivation impairs blood-feeding propensity, with a significantly lesser proportion of mosquitoes able to blood feed in the sleep deprived group (about 54% reduction) in comparison with control (Wilcoxon rank sum test: W = 58.5; *p* < 0.01; Figure 4C).

Overall, host landing is significantly suppressed by sleep deprivation in both lab and field conditions and sleep deprivation induced a reduction in blood feeding during the periods when *Ae. aegypti* females are typically active.

## Discussion

Our studies establish the occurrence of sleep-like states in mosquitoes including *Ae. aegypti*, *Cx. pipiens* and *An. stephensi* based on some of the conventional behavioral features described in other insect systems. These consist of a consolidated period of inactivity/immobility in a particular phase of the circadian day, postural differences between active (awake) state and putative sleep state, and the occurrence of sleep recovery following sleep disruption. Lastly, the influence of sleep deprivation on mosquito biology and their role in disease transmission was established by identifying that a reduced arousal while in sleep states when a host is present and host landing and blood feeding patterns can be altered by sleep deprivation.

Sleeping arthropods assume obvious sleep postures. For example, antennal positions are associated with sleep in *A. mellifera*, where the scapes are positioned almost horizontally close to the head surface, and the pedicels with their flagella assume a vertical position during the night, which are different during locomotor activity in the subjective day [20]. In the same insect system, small swaying movements of the antennae are associated with the resting state [20]. In the nocturnal cockroach, *Blaberus giganteus*, raised body posture and antennal movements are predominant in the dark period, while rest during the day is associated with the body and the antennae touching the substrate [41]. In *D. melanogaster*, individual flies move away from their food source and take up a prone position prior to resting [18]. Respiratory abdominal pumping and small sporadic proboscis extension/retraction are the only movements that occur during the sleep-like state in these flies [18]. Evidence for postural differences between active and sleep-like states in mosquitoes was successfully established in our study for three mosquito species, with whole body orientation and most importantly hind leg angle providing significant distinctions between these states. This is the first study in insects where the orientation of the insect leg is a feature distinguishing sleep-like condition from the active state. Interestingly, a subtle difference was seen between the culicine species (*Ae. aegypti* and *Cx. pipiens*) and the anopheline species (*An. stephensi*). In the latter species, leg angle was not strong enough to show conspicuous difference between the active and sleep-like states. Unlike what is found in the culicines, during non-flight activity, adult *Anopheles* mosquito typically has its abdomen pointing away from the substrate, thereby forming an angle of 30-45° with the substrate (or resting surface) [42]. Legs of anophelines are generally longer than those of culicines [42]; these might explain the weak difference between active and sleep-like states based only on leg orientation.

Historical observations of biting/feeding and resting behavior in the field have shown that these occur at different periods of the day in mosquito species, being modulated by circadian rhythms [43, 44]. *Aedes* mosquitoes feed mostly and are active during the day, while *Cx. pipiens* and *An. stephensi* have increased feeding activity during the twilight and night respectively; these differences in feeding and resting time matched our laboratory observations in this study [28,38,40]. Furthermore, the reduction in arousal when in the sleep states we observed in *Ae. aegypti* and *Cx. pipiens* could be a contributing factor in why these mosquitoes do not observe feeding, even when a host is present, during the night and day, respectively. In *Drosophila-*based studies where the flies were subjected to 12hr: 12hr photophase/scotophase, prolonged periods of rest were observed in the dark period, similar to what we observed in *Ae. aegypti* [19,37,45]. The most significant difference between these studies and this current one is the duration of immobility used to establish sleep. While a 5-min period of inactivity was sufficient to define sleep in *Drosophila*, a period of no activity for at least 120 minutes was used in our study for mosquitoes based on postural and arousal observations. The strong preference for rest in the photophase for *An. stephensi* and *Cx. pipiens* is similar to what was reported in two cockroach species; *Leucophaea maderae* and *B. giganteus* [3, 41]. Furthermore, the difference in circadian timing of sleep-like states (low activity) observed among the different mosquito species in this study is not surprising, as there are reports in other studies of differences in several aspects of activity/rest rhythm among closely related species, but these studies are limited to a few comparisons in fruit flies and wasps [46, 47]. Sleep rebound is an important hallmark of sleep, where there is an increase in sleep following sleep disruption in the phase an individual normally sleeps i.e. a homeostatic regulation of sleep [15]. This phenomenon has been confirmed in different arthropods including scorpions [48], cockroaches [3, 41], honey bees [49], and fruit flies [18, 19], and also in other non-arthropod systems [1,2,50]. In the present study, we observed an increase in sleep amount in the subsequent phase as a result of nighttime and daytime sleep deprivation in *Ae. aegypti* and *An. stephensi*, respectively. As expected, daytime sleep deprivation did not induce sleep rebound in *Ae. aegypti*, similar to the result for daytime sleep deprivation in a *Drosophila*-based study [19]. This indicates a compensatory increase in sleep following nighttime sleep deprivation in *Ae. aegypti* was not driven by increased activity but by sleep loss.

Sleep deprivation impacts a diverse range of biological processes in animals including cognition, metabolism, alertness, reproduction, and immunity [51, 52]. In honey bees, foraging efficiency of nestmates is affected due to the negative effect of sleep deprivation on waggle dance signaling [11]. Short- and long-term memory are both disrupted by nighttime sleep deprivation in *Drosophila* [12, 13], but adequate sleep tends to facilitate memory and learning improvement [53, 54]. In another study, sleep deprivation in *Drosophila* was reported to suppress aggressive behaviors, with a serious implication on reproductive fitness [55]. Importantly, a strong link between sleep and immune function have been established in *Drosophila* [56], and studies have shown that reduced sleep leads to increased resistance to bacterial infection and a major category of genes that increased expression due to sleep deprivation is involved in immune function [57, 58]. These results are particularly interesting for our study system because circadian rhythms modulate immune response [59], and immunity is one of the main factors that influence disease transmission in mosquitoes [26]. Although our study was solely behavioral elucidation of sleep, and we did not consider the influence of sleep deprivation on immune response, however, we were able to show the potential effect of sleep disruption on mosquito’s vectorial capacity by measuring host landing, blood-feeding propensity and arousal when host is present, which are critical to obtain a blood meal and transmit pathogens [60]. Laboratory and field mesocosm experiments revealed that *Ae. aegypti* mosquitoes had significantly reduced response to a host mimic after nighttime sleep deprivation. Based on studies in other systems, sleep deprived *Ae. aegypti* are sleeping more during the day to recover their lost sleep from the previous night, thereby displaying an increased arousal threshold to host stimulation - an important hallmark of sleep [15]. The successful transmission of diseases by mosquitoes is heavily reliant on a pathogen-carrying mosquito encountering a host at a specific time that matches, and eventually introducing the infective stage of the pathogen to the host during feeding [61]. We predicted that sleep deprivation will affect disease transmission, since blood-feeding propensity was also significantly impaired in our study. Furthermore, the acquisition of a specific pathogen requires the vector feeds at a specific time when stages are present in the blood that can establish within the vector [62, 63]. Hence, altered host landing and blood feeding in mosquitoes due to sleep deprivation could change the dynamics between host, pathogens, and disease vector. These interacting aspects indicate there is an urgent need to investigate the influence of sleep deprivation on other components of vectorial capacity as this would improve current disease modeling and vector control strategies.

## Materials and Methods

### Mosquito husbandry

Three mosquito species were used; *Aedes aegypti*, *Culex pipiens*, and *Anopheles stephensi*. *Culex pipiens* colonies used for this study were originally collected in 2015 from Columbus, OH, and supplemented with field-collected individuals every two to three years (Buckeye strain), while those of *Ae. aegypti* and *An. stephensi* were acquired from Benzon Research (Carlisle, PA, USA) and BEI Resources (*Ae. aegypti,* Rockefeller strain, MR4-735; *An. stephensi*) for postural analysis. Mosquito colonies were maintained in the laboratory at the University of Cincinnati at 25°C, 80% relative humidity (RH) under a 15hr: 9hr light/dark (L/D) cycle with access to water and 10% sucrose *ad libitum* and at Virginia Tech under the same conditions for the postural analysis. Mosquito eggs were produced from 4-5 weeks old females through artificial feeding (Hemotek, Blackburn, United Kingdom) with chicken or rabbit blood (Pel-Freez Biologicals, Rogers, AZ, USA). Upon egg hatching, larvae were separated into 18 cm x 25 cm x 5 cm containers (at a density of 250 individuals per container) and were fed finely ground fish food (Tetramin, Melle, Germany). For the experiments, pupae were collected and maintained in an incubator at 24°C, 70-75% RH, under a 12hr:12hr L/D cycle until adult emergence. Adult mosquitoes that emerged were provided with access to water and 10% sucrose *ad libitum*. Unless otherwise stated, all adult female mosquitoes used for the laboratory-based experiments were aged 5-8 days post-ecdysis. However, adult female mosquitoes (12-17 days old) were collected directly from the maintained laboratory colonies for the field-based experiments. As the experimenters represent potential blood-host to the mosquitoes in all experiments, studies were conducted in isolated experimental rooms and incubators to eliminate potential disturbances from the experimenter. Remote computer access and automated data collection were used to prevent exposure to host-based factors.

### Posture analysis

#### Quantification of postural changes associated with prolonged immobility

To quantify body postures associated with putative sleep states, groups of 20, 5-7 day old adult females of *Ae. aegypti*, *Cx. pipiens*, and *An. stephensi* were enclosed within acrylic containers (16 Oz mosquito breeder; BioQuip, Rancho Dominguez, CA, USA) covered by a fabric mesh at the top. Containers were positioned within the field of view of an infrared camera (PointGrey Firefly MV FMVU-03MTC, FLIR, Wilsonville, OR, USA) connected to a computer. After the experimenter left the room, mosquitoes were left unperturbed for 2 hours to allow acclimatation to the experimental environment. Then, the experimenter remote-accessed the computer and pictures of individual mosquitoes were taken during a one hour window. Only mosquitoes that were landed perpendicular to the focal plane of the camera, with their legs clearly visible were conserved for the analysis (*Ae. aegypti:* n = 22; *Cx. pipiens*: n = 41; *An. stephensi*: n = 17). All experiments were conducted on sugar fed but never blood fed females during the last 2 hours of the photophase. Depending on whether the focal mosquito was seen moving its appendages (*e.g.*, grooming, moving of the legs), it was either classified as “active” or “at rest”. Saved images were imported in ImageJ (National Institutes of Health, USA) where the hind leg angle relative to the mosquito’s main body axis, the body angle relative to the substrate, the elevation of the hind leg relative to the substrate, and the elevation of the thorax relative to the substrate were measured. All length measurements (in pixels) were normalized to the length of the mosquito’s body, from tip of the abdomen to the top of the thorax. Repeated measurements of the same image showed a tracking error of 2.41 pixels for lengths, which represents a fraction of the thickness of the hind legs, and an error of 0.61 degrees for angles. A Principal Component Analysis (PCA) was conducted in R version 3.6.3, and an ANOSIM (package *vegan* version 2.5-6) was performed to test for the dissimilarity between species, as well as between “active” and “at rest” mosquitoes.

#### Time course analysis of body postures

Adult females of each species were individualized in plastic *Drosophila* tubes (25mm x 95mm, Genesee Scientific, San Diego, CA, USA) and, for each replicate (n =3), 20 tubes of females of the same species were positioned horizontally, in the field of view of a video camera (C920, Logitech, Lausanne, Switzerland). Every 10 minutes for the last 3 hours of the photophase and for the first 3 hours of the scotophase, the posture of each individual was recorded and classified as ‘active’ or ‘at rest’ based on the angle of the hind legs relative to the main body axis. Analysis of the data was performed in R.

#### Basic rest-activity rhythms

The rest-activity rhythms of the three mosquito species were quantified using a Locomotor Activity Monitor 25 (LAM25) system (TriKinetics Inc., Waltham, MA, USA) and the DAMSystem3 Data Collection Software (TriKinetics). Originally, these systems were developed for *Drosophila* but recently have been utilized to measure the activity levels of several blood-feeding arthropods, including mosquitoes [64–66]. Individual mosquitoes were placed in 25 x 150 mm clear glass tubes with access to water and 10% sucrose provided *ad libitum*. These tubes were placed horizontally in the LAM25 system which allows the simultaneous recording of 32 mosquitoes in an “8 x 4” horizontal by vertical matrix during a single trial. The entire set-up was held in a light-proof low-temperature incubator supplied with its own lighting system at 24°C, 70-75% RH, under a 11hr:11hr L/D cycle (with 1hr dawn and 1hr dusk transitions). After the acclimation of the mosquitoes for 2 days, activity level was recorded as the number of times (in a minute) a mosquito crosses an infrared beam of the LAM25 in the middle of the locomotor tube. Data collected with the DAMSystem3 for a duration of 5 days (with the removal of mosquitoes that were not alive till the end of the assay) were analyzed using the Rethomics platform in R with its associated packages such as *behavr*, *ggetho*, *damr* and *sleepr* [67].

#### Sleep deprivation assay

Following the acclimation of the mosquitoes for 2 days and the establishment of a 24-hour baseline day in the LAM25 system, sleep deprivation was conducted in the specific phase of interest. Sleep deprivation was achieved in the mosquitoes through the delivery of vibration stimuli (vibration amplitude = 3G) using a Multi-Tube Vortex Mixer (Ohaus, Parsippany, NJ, USA) attached to the LAM25 system. In the diurnal *Ae. aegypti* mosquitoes, three different sleep deprivation protocols were conducted based on modifications from a *Drosophila* study [55]: 12hr nighttime deprivation (12NTD), 4hr nighttime deprivation (4NTD) and 12hr daytime deprivation (DTD). Whereas in the nocturnal *An. stephensi* mosquitoes, only DTD was conducted. To accomplish 12NTD, a sequence of vibration pulses lasting 1 minute, followed by 5 minutes of rest between pulses was programmed for the entire scotophase subsequent to the baseline day (see Figure S1A). In 4NTD, vibration pulses lasted for 1 minute followed by 1 minute of rest between pulses in the first 4 hours of the night (Zeitgeber time 12 - 16) following the baseline day. This was done in such a way that the total number of vibration pulses obtainable in 12NTD was delivered in a short time frame (see Figure S1B). DTD in *Ae. aegypti* and *An. stephensi* was conducted similarly to 12NTD, the only difference is that DTD was accomplished during the photophase that succeeds the baseline day (see Figures S1C and S1D).

To calculate sleep loss, we used the mean difference of the sleep amounts in the scotophase (12NTD and 4NTD) or photophase (DTD) preceding the deprivation and that of the scotophase (12NTD and 4NTD) or photophase (DTD) during the deprivation. For the calculation of sleep gain, we used the mean difference of the sleep amounts in the photophase (12NTD and 4NTD) or scotophase (DTD) after the deprivation and that of the photophase (12NTD and 4NTD) or scotophase (DTD) before the deprivation.

#### Host landing and blood feeding assays

Host landing following 4 hours post-sleep deprivation (PSD) was assessed in *Ae. aegypti* mosquitoes both in laboratory and field conditions. In the lab-based study, sleep deprivation protocol was similar to 12NTD (described under “Sleep deprivation assay”). The only difference was that a “17.5 cm x 17.5 cm x 17.5 cm” knitted mesh-nylon cage (BioQuip) housing mosquitoes (10 per replicate) was attached to the Multi-Tube Vortex Mixer to achieve sleep loss, with the entire set-up held in a room isolated from host cues (26 +/- 1°C, 75 ± 5% RH, and 12hr:12hr L/D cycle). In the field-based experiment, adult female mosquitoes were released into similar cages described earlier (10 mosquitoes per replicate), which were then placed into “47.5 cm x 47.5 cm x 47.5” knitted mesh-nylon cages (BioQuip). To achieve bulk sleep deprivation, the entire set-up was situated in a city environment, where high activity occurs. The control set-up was also placed in the same environment as the sleep-deprived counterpart, but located in a secluded area that experiences significantly reduced disturbances. To determine the number of mosquitoes that landed on a host mimic, we used techniques adapted from previous studies [68, 69]. A host mimic (Hemotek feeder) filled with a mixture of water and 100µl artificial eccrine perspiration (Pickering Laboratories, Mountain View, CA, USA) heated to 37°C was covered three times with parafilm and placed on top of the experimental cage. Incidental contact was distinguished from foraging contact by using mosquitoes that landed and remained for at least 5 seconds on the feeder. By recording using a video camera (7 White, GoPro, San Mateo, CA, USA), the number of mosquitoes that made foraging contact was counted in the lab experiment at 10 mins, 20 mins, 40 mins and 60 mins after the artificial host was turned on. In the field experiment, this was counted only after 5 mins, 10 mins and 15 mins. Result was expressed as a proportion of the total mosquitoes that remained alive at the end of the experiment, and compared with the control group.

To assess the influence of sleep deprivation on blood-feeding propensity in mosquitoes, Adult female *Ae*. *aegypti* were exposed to the legs of a volunteer human host for 5 minutes after 4 hours PSD (Approved by the University of Cincinnati IR 2021-0971). The set-up and sleep deprivation protocol in this experiment were similar to the lab-based host landing assay described above. The number of mosquitoes that successfully blood fed (shown by engorged abdomen) in the sleep-deprived group was compared to that of control (non-sleep deprived group), and expressed as a proportion of the total mosquitoes that stayed alive throughout the duration of the assay.

#### Reduction in host responsiveness

To determine if prolonged sleep-like states reduced the response of mosquitoes, we performed basic host cue response studies on two species (*Ae*. *aegypti* and *Cx*. *pipiens*). Mosquitoes were observed through video and after 0, 30, 60, 120, and 240 minutes of inactivity, the experimenter entered the room and exhaled on the cage to provide a host cue. The number of mosquitoes that took flight within thirty seconds following exposure to experimenter breath was used as a proxy for host response. Each time point was conducted on 8-14 mosquitoes for each species.

#### Quantification and statistical analysis

Experimental replicates utilized for the study are distinct samples and biologically independent. Sample sizes for the different experiments are mentioned in the methods or in the associated figure legend. Statistical tests, and significance between groups are detailed within each figure and/or in the figure legend. All analyses were done in R version 3.6.3.

## Supplemental materials Figures

**Figure S1.**
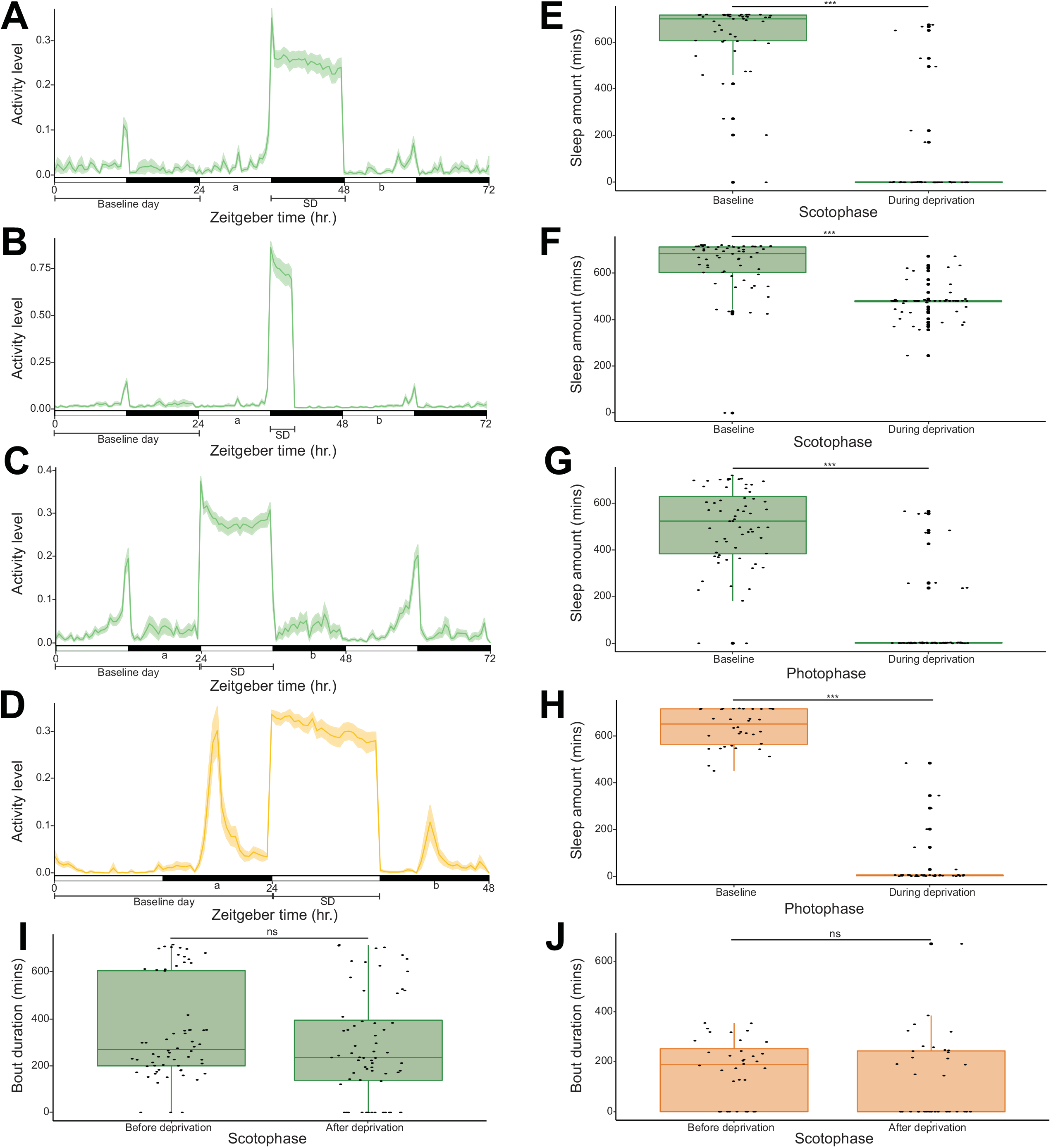
Experimental design and activity profile of (A) 12hr nighttime sleep deprivation experiment in *Aedes aegypti* (n = 48), (B) 4hr nighttime sleep deprivation experiment in *Aedes aegypti* (n = 59), (C) 12hr daytime sleep deprivation experiment in *Aedes aegypti* (n = 64) and (D) 12hr daytime sleep deprivation experiment in *Anopheles stephensi* (n = 36). The y axis represents the mean beam crosses made by all the mosquitoes, and the x axis represents the Zeitgeber time. The solid lines and the shaded areas show population means and their 95% bootstrap confidence interval, respectively. White and black horizontal bars represent the photophase and scotophase, respectively. ‘a’ denotes the phase before sleep deprivation and ‘b” denotes the phase after sleep deprivation. Comparison of sleep amounts between baseline and during sleep deprivation in (E) 12hr nighttime sleep deprivation experiment in *Aedes aegypti* (n = 48), (F) 4hr nighttime sleep deprivation experiment in *Aedes aegypti* (n = 59), (G) 12hr daytime sleep deprivation experiment in *Aedes aegypti* (n = 64) and (H) 12hr daytime sleep deprivation experiment in *Anopheles stephensi* (n = 36). Comparison of average bout durations before and after sleep deprivation in (I) 12hr daytime sleep deprivation experiment in *Aedes aegypti* (n = 64, individuals with zero values were included) and (J) 12hr daytime sleep deprivation experiment in *Anopheles stephensi* (n = 36, individuals with zero values were included). Test of significant difference between groups was carried out using wilcoxon signed rank test (ns = not significant, *** = *p* < 0.001).

## Videos

Supplemental - Videos of host landing between control and sleep-deprived mosquitoes.

Available upon request.

## Acknowledgements

We thank Diane Eilerts for fruitful discussions about the project, and Zhijian Jake Tu for providing females of the Indian strain of *Anopheles stephensi*, a representative of the type form. The following reagent was obtained through BEI Resources, NIAID, NIH: *Aedes aegypti*, Strain LVP-IB12, Eggs, MRA-735, contributed by David W. Severson.

